# Using low pass whole metagenome sequencing for gut microbiome profiling in an Argentine urban population

**DOI:** 10.1101/2025.07.31.667939

**Authors:** Milagros Trotta, Cristian Rohr, Guadalupe Benavidez, Martin Vazquez

## Abstract

Modern urban diets are linked to gut microbiome dysbiosis and increased risk of chronic disease. While 16S rRNA sequencing is widely used, low-pass whole metagenome sequencing (WMS) offers superior insight into microbial functional capacity and community structures. In this study, we utilized low-pass WMS to characterize the gut microbiome of an Argentine urban population. We compared a health-screened ‘Reference’ cohort (n=94) with an unscreened ‘Average’ urban cohort (n=527), for which we analyzed taxonomic structure, microbial diversity, and predicted metabolic potential.

The ‘Average’ cohort displayed significantly lower alpha-diversity, primarily driven by reduced community evenness rather than richness. Functionally, this cohort showed a significantly diminished predicted capacity for synthesizing beneficial short-chain fatty acids (SCFAs) and essential vitamins B9 and B12. A longitudinal analysis of a subgroup (n=9) undertaking lifestyle modifications demonstrated that while microbial richness and SCFA production could be significantly improved, responses were highly individualized.

These results reveal key functional deficits in the gut microbiome of a typical urban population, reflecting a mismatch with ancestral dietary patterns. The high inter-individual variability in response to changes in dietary patterns challenges one-size-fits-all dietary recommendations and highlights the need for personalized nutritional strategies informed by functional microbiome data.

## INTRODUCTION

The human gut microbiome, a complex and dynamic ecosystem comprising trillions of microorganisms, has emerged as a critical player in host health, influencing a broad spectrum of physiological processes including nutrient metabolism, immune system development, and protection against pathogens [1]. Dysbiosis, or imbalances in the gut microbial community, has been increasingly implicated in the pathogenesis of numerous chronic diseases, including inflammatory bowel disease (IBD) [2], obesity [3], type 2 diabetes [4], cardiovascular disease [5], and even neurological disorders [6]. This has fueled a surge of interest in gut microbiome profiling as a potential tool for disease diagnosis, risk stratification, and the development of targeted interventions, particularly through nutritional modulation. Dietary interventions, such as prebiotics, probiotics, and personalized diets, are being actively explored as strategies to reshape the gut microbiome and mitigate disease risk [7], [8].

Historically, human gut microbial communities were shaped by diets rich in fiber and diverse plant matter, reflecting an intimate relationship with the natural environment. However, the rapid industrialization and globalization of food production have dramatically altered dietary patterns worldwide, particularly with the increased consumption of ultra-processed foods (UPFs). These UPFs, often characterized by high sugar, salt, and fat content, along with a paucity of fiber and essential nutrients, are now a dominant feature of modern diets, especially in urban environments [9].

Essentially, early hominids primarily consumed plants, with meat being a more limited, opportunistic component of their sustenance. Key innovations, such as the control of fire and the subsequent advent of agriculture, significantly altered food preparation and availability. These developments led to increased food energy accessibility but also, notably, a reduction in nutritional intake diversity [10]. The profound divergence between these ancestral genetic predispositions and contemporary dietary environments sets the stage for what can be described as an “**evolutionary mismatch**” in modern nutrition. This fundamental incongruity provides a powerful, overarching framework for understanding the root causes of many chronic diseases prevalent in industrialized societies. These diseases are not merely random occurrences or solely the result of individual lifestyle choices, but rather a direct consequence of a deep biological-environmental disconnect, where human bodies, and specifically their microbiomes, are ill-equipped to optimally process and respond to the modern food landscape [10]. This reframes the public health challenge from simply managing symptoms to addressing a deep-seated evolutionary incongruity.

The shift towards industrialized diets and UPF consumption has been linked to significant disruptions in gut microbiome composition and function [11]. Studies have demonstrated that diets high in UPFs can lead to reduced microbial diversity, altered metabolic activity, and increased susceptibility to inflammatory conditions [11]. Specific consequences include a decrease in beneficial bacterial taxa, such as *Bifidobacterium* and *Lactobacillus*, and an increase in opportunistic pathogens and bacteria associated with inflammation, potentially contributing to the rise of chronic diseases like obesity, type 2 diabetes, and inflammatory bowel disease [12]. Furthermore, the additives and emulsifiers commonly found in UPFs can directly impact the gut barrier, leading to increased intestinal permeability and systemic inflammation. Understanding the impact of industrialization and UPF consumption on the gut microbiome is particularly important in rapidly urbanizing regions of the world [13].

Traditionally, microbiome profiling has relied heavily on 16S rRNA gene sequencing, a culture-independent method that targets a highly conserved region of the bacterial genome. While 16S rRNA sequencing offers a cost-effective and relatively simple approach for identifying and quantifying bacterial taxa within a sample, it suffers from several limitations. Firstly, its taxonomic resolution is often limited to the genus level, hindering the identification of specific strains. Secondly, 16S rRNA sequencing provides limited information about the functional potential of the microbiome, failing to capture the metabolic pathways and enzymatic capabilities encoded by the microbial community [14]. In contrast, whole metagenome sequencing (WMS) offers a more comprehensive and nuanced approach to microbiome profiling. WMS involves sequencing the entire DNA content of a microbial sample, providing access to the genetic blueprint of all microorganisms present, including bacteria, archaea, viruses, and fungi. This allows for taxonomic profiling at a higher resolution, often down to the species and even strain level, as well as the identification of functional genes, metabolic pathways, and antibiotic resistance genes. Furthermore, WMS enables the investigation of microbial interactions and the construction of metabolic networks, providing a deeper understanding of the ecological dynamics within the gut. While WMS offers significant advantages over 16S rRNA sequencing, it is also more computationally intensive and generally more expensive [15], [16], [17].

Despite the increased cost, the superior resolution and functional insights provided by WMS are increasingly recognized as essential for translating microbiome research into clinical applications. Indeed, recent microbiome testing guidelines emphasize the importance of selecting appropriate sequencing methods based on the specific research question and the desired level of detail [15]. The choice between 16S rRNA sequencing and WMS should be carefully considered, considering the trade-offs between cost, resolution, and the type of information required [14]. For studies aiming to identify specific microbial species associated with disease or to elucidate the functional mechanisms underlying microbiome-host interactions, WMS is generally the preferred approach [16]. Low-pass WMS, a cost-effective alternative to deep sequencing, can still provide sufficient coverage for accurate taxonomic and functional profiling, particularly in well-defined communities [18].

Given the growing importance of gut microbiome profiling in understanding and addressing chronic diseases and the limitations of traditional 16S rRNA sequencing, this study aimed to reconstruct our reference-controlled dataset for Argentine urban population using low-pass whole metagenome sequencing. We then used the reference-controlled dataset to compare against a cohort of an average (not controlled) urban population from our disease prevention program. We aim to define the gut microbial profile and metabolic capacity within this population looking for patterns that can be translated into effective nutritional interventions, ultimately mitigating the adverse health effects associated with modern, industrialized diets.

## MATERIALS AND METHODS

### Study cohorts

The Argentinian reference cohort was previously described [[19]. Briefly, 200 volunteers were screened in four different cities of Argentina, after an extensive health and habits questionnaire and blood, urine and metabolome tests, 94 volunteers were selected for the reference-controlled program and stool samples were taken to analyze gut microbiome. This cohort was selected based on strict criteria for healthy habits (Reference cohort) since they passed the blood and urine screening for all major clinical biomarkers using standard clinical laboratory analysis and metabolome [[19]. The average age of this cohort was 32 years with a standard deviation of 8 years, 43.6% of the participants were male and average body mass index was 24.2 ± 3.2 kg/m^2.^ Participants represent typical middle-class of large Argentine cities in similar socioeconomic status (SES). 90% of the participants reported consuming a balanced varied diet, including all major types of food like meat, vegetables, grains, fruits and dairy, with no changes in their diets for at least six months.

Another cohort of 527 participants was anonymously selected from Heritas′ wellness and prevention program. Participants agreed to participate anonymously in research studies upon acceptance of the terms and conditions of the program. This cohort was not strictly controlled for healthy habits and is considered a standard average population living in industrialized cities (Average cohort). The average age of this cohort was 45 years with a standard deviation of 13 years, 47.4% of the participants were male and average body mass index was 24.2 ± 4.6 kg/m^2^. Participants represent typical middle-class of large Argentine cities in similar socioeconomic status (SES). In fact, SES was very similar in both cohorts by design to avoid it as a confounding factor. Regarding dietary habits, 53% reported consuming a varied diet, and an analysis of specific food consumption revealed that most participants reported consuming an average of up to two daily portions of vegetables, fruits, and legumes, while consumption of whole grains and nuts was less frequent, and nearly half of the cohort reported consuming yogurt daily.

A small random subset of the Average cohort, nine participants (US01-US09), were enrolled for a longitudinal analysis after lifestyle adjustments. Our inclusion criteria required individuals to have completed two gut microbiome assessments (T0 and T1) separated by three months, and to have a self-reported dietary and lifestyle modifications questionnaire completed (Suppl Material File 1).

### Gut Microbiome Analysis

Fecal samples were collected by volunteers or program users using DNA/RNA Shield™ Fecal Collection Tubes (Zymo Research). DNA extraction from samples was performed using the QIAamp Fast DNA Stool Mini Kit (QIAGEN). The extracted DNA was quantified using the Quant-iT PicoGreen dsDNA Assay Kit (Invitrogen). DNA libraries were prepared with the Nextera XT DNA Sample Prep Kit (Illumina). Stool samples from both cohorts underwent low-pass whole metagenomic sequencing (WMS), targeting a minimum of 1.3 million reads per sample. The sequencing was performed on a NextSeq 550 sequencer following standard Illumina protocols. An internal control was included in all sequencing runs to monitor and ensure technical reproducibility.

Bioinformatic analyses were performed using SqueezeMeta v1.4 [20] run in co-assembly mode with default parameters to identify microbiome species and functionality, requiring a minimum of 500,000 mapped reads per sample and a mapped reads-to-total reads ratio greater than 0.7. Alpha diversity was calculated as Shannon index, while richness using the observed taxon and evenness as alpha diversity / log(richness).To assign Short-Chain Fatty Acids (SCFAs), Trimethylamine (TMA), and the synthesis of vitamins B1 and B7, Vitamin B12, Vitamin B9 and Vitamin K2 to the metagenomic samples, KEGG (Kyoto Encyclopedia of Genes and Genomes; release 103) identifiers were used. The KEGGs IDs used for each metabolite are detailed in Supplementary table 1. The metabolic production capacity for each mentioned metabolite was calculated as the sum of the FPM (Fragments Per Million Reads) values for each detected KEGG entry, ensuring sample normalization.

### Statistics Analysis

Data obtained from species and production capacity was analyzed using matplotlib and SciPy phyton libraries. Statistical significance was defined as a p-value < 0.05. The homoscedasticity of the samples was considered to decide whether to perform parametric or non-parametric tests.

## RESULTS

### Community structures in the reconstructed reference-controlled dataset of the Argentine urban population using low-pass WMS

We previously described a reference-controlled gut microbiome dataset for a healthy urban cohort in Argentina comprised of 172 individuals [19]. The dataset was constructed using NGS of 16S rRNA v3-V4 regions. We reconstructed the dataset using low-pass WMS technology from 98 DNA samples from the same cohort. Clustering analysis of the low-pass WMS data identified three distinct microbial community types (Clusters 1, 2, and 3) based on their characteristic taxonomic composition at the genus level (Figure 1A, Table 1).

**Figure 1.**
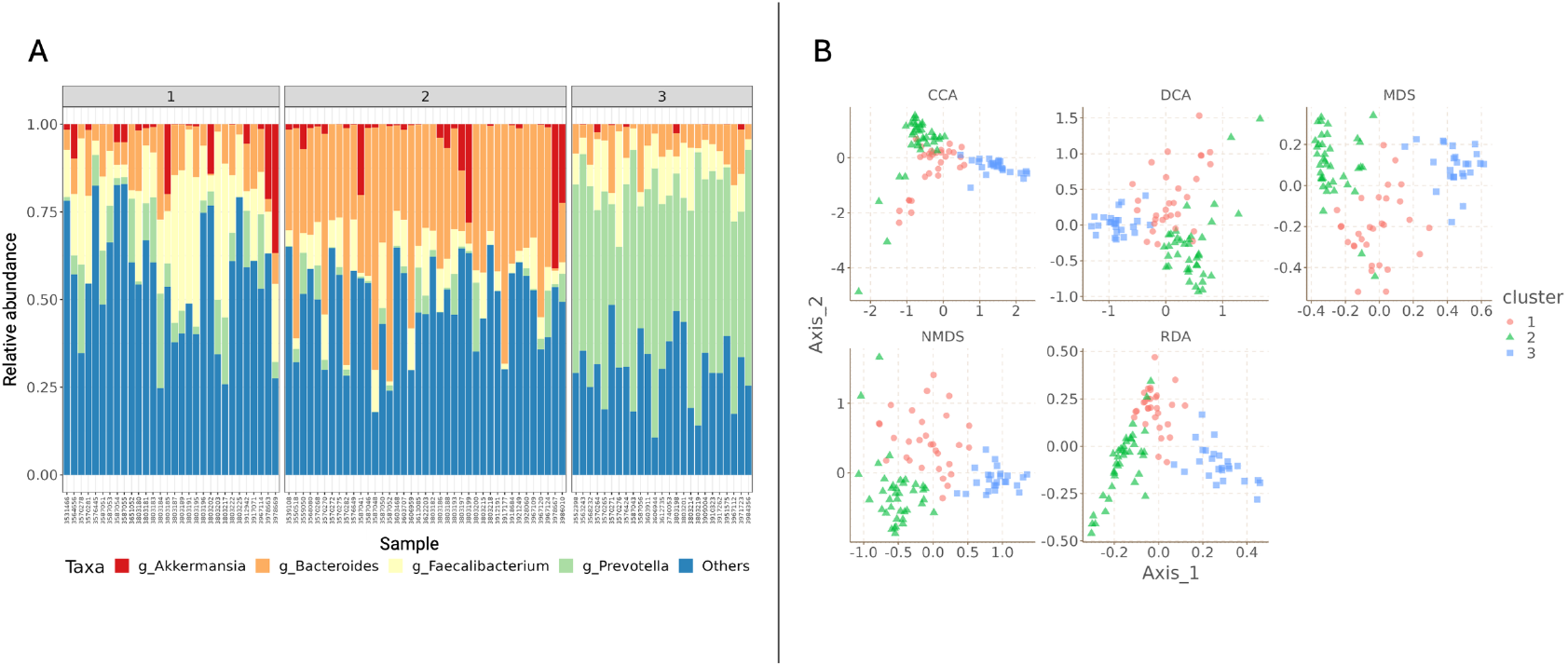
(A) Clustering analysis of the low-pass WMS data from the reference group. (B) Multidimensional space visualization using different multivariate ordination methods. Samples from Cluster 1 (red), Cluster 2 (green), and Cluster 3 (blue) consistently group together and show distinct separation from samples of other clusters, with minimal overlap

**Table 1.** Overview of the clustering analysis of the low-pass WMS data from the reference group, indicating key taxonomic characteristics of the dominant genera in each cluster.

| Cluster ID | Dominant Genera | Key Taxonomic Characteristics |
| --- | --- | --- |
| **Cluster 1** | *Faecalibacterium* | High relative abundance of *Faecalibacterium*; *Bacteroides* and *Prevotella* are present but not dominant. |
| **Cluster 2** | *Bacteroides* | High relative abundance of *Bacteroides*; comparatively lower *Prevotella* and *Faecalibacterium*. |
| **Cluster 3** | *Prevotella* | High relative abundance of *Prevotella*; markedly reduced *Bacteroides*. |

A notable aspect revealed by the low-pass WMS data, particularly when considering the context of our previous 16S rRNA V3-V4 gene sequencing work, is the role of *Akkermansia* in the clustering structure. While *Akkermansia* defined a cluster per se in our previous work [19], now appeared more distributed in all clusters, particularly 1 and 2, and not defining one (Table 1, Figure 1A).

To visualize and confirm the distinctness of the identified microbial clusters in a multi-dimensional space, a comprehensive array of multivariate ordination methods was employed (Figure 1B). These methods included Principal Coordinate Analysis (PCoA) utilizing various distance metrics such as Jaccard (presence/absence based), Bray-Curtis (abundance-based), Unifrac (phylogenetic, unweighted), and Weighted Unifrac (phylogenetic, weighted by abundance). Additionally, Canonical Correspondence Analysis (CCA), Detrended Correspondence Analysis (DCA), Non-metric Multidimensional Scaling (NMDS), and Redundancy Analysis (RDA) were performed.

A consistent finding across all six ordination plots is the clear and robust separation of samples belonging to the three identified clusters (Figure 1B).

We performed a direct comparison between 16S rRNA V3-V4 sequencing and low-pass WMS by randomly selecting four different samples from the dataset pools. Results indicated that 16S overestimated Firmicutes and Proteobacteria while underestimating Bacteriodetes at the phylum level. On the contrary, low-pass WMS reproducible showed the opposite situation and persistently identified more diverse phylums, including “unclassified” categories as expected for this type of technology (Figure 2A). This discrepancy was magnified at the genus level (Figure 2B). Low-pass WMS showed greater diversity of genus in comparison with 16S. Moreover, “Unknown”/”Uncultured” categories appeared in the 16S dataset showing a proportion of reads that cannot be mapped. On the contrary, a greater number of reads in low-pass WMS were mapped to unclassified Bacteroidales and Clostridiales that 16S failed to map. Overall, as expected, low-pass WMS offered a greater resolution on gut microbiome profiling over 16S rRNA V3-V4. This highlights a significant advantage of transitioning our reference dataset from 16S to low-pass WMS, as WMS offers a more complete profile of the microbial community, including access to the predicted metabolic capacity.

**Figure 2.**
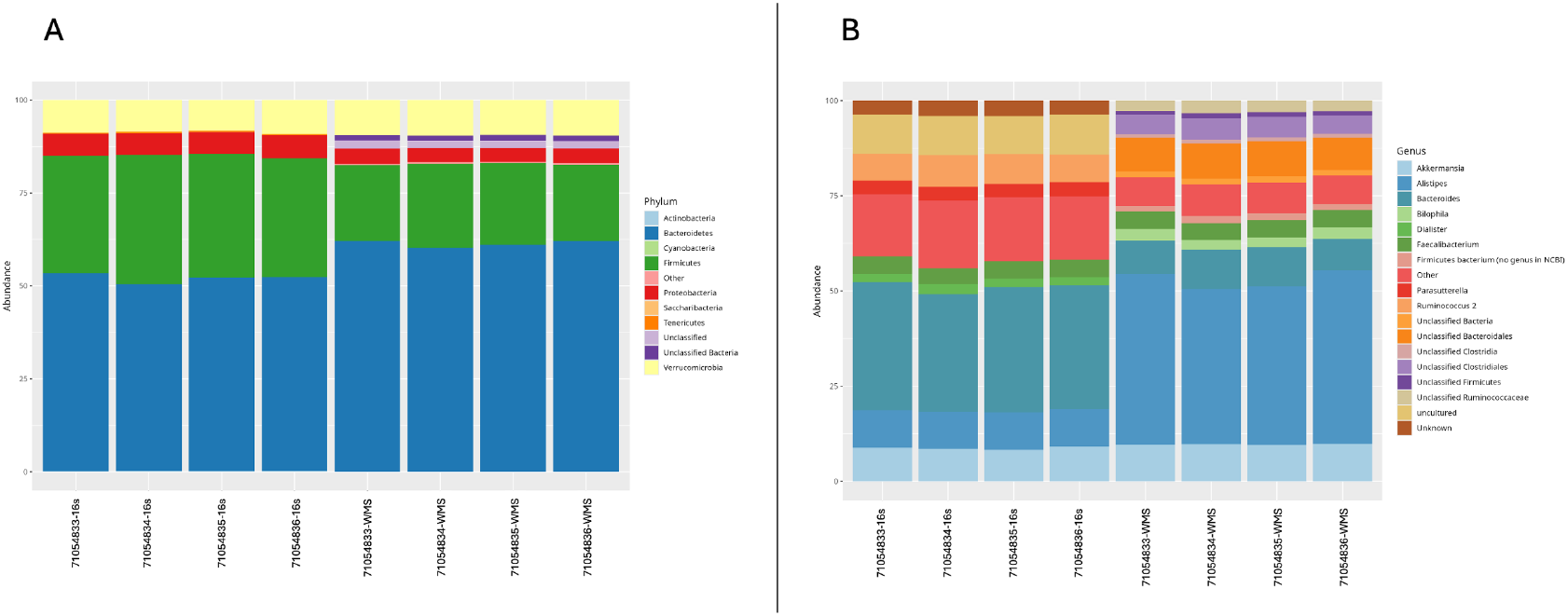
16S rRNA V3-V4 sequencing vs low-pass WMS. (A) Phylum level comparison. (B) Genus level comparison. Four samples were randomly selected from the dataset and compared. Duplicated samples for low-pass WMS came from different Illumina library preparation methods.

### Gut microbiome profiles in the average urban cohort vs reference cohort

Having reconstructed the reference cohort using low-pass WMS, we analyzed and compared other cohorts using the same technology. To characterize the gut microbiome of the Argentinian average urban population, we selected 527 participant samples from Heritas′ wellness and prevention program to compile an average urban cohort. Participants were not health-screened and considered average middle-class population from large Argentine cities, as described in Materials and Methods section. Then, we analyzed differences in the community structure and reconstructed metabolic potential of key indicators between average and reference cohorts.

We analyzed three common indicators of microbial community structure: Shannon alpha-diversity, genus richness, and genus evenness, comparing them to those of the reference cohort. Alpha-diversity (Figure 3A), as measured by the Shannon index, was significantly higher in the reference cohort (mean: 2.98 ± 0.032) compared to the general population cohort (mean: 2.71 ± 0.016; p < 0.001). In contrast, no significant differences were observed in genus richness (Figure 3B) between the cohorts (reference mean: 334.34 ± 4.66, general population mean: 344.61± 3.01; p > 0.05), indicating that the number of total genera was comparable. However, genus evenness (Figure 3C) was significantly greater in the reference cohort (reference mean: 0.51 ± 0.005, general population mean: 0.46± 0.002; p < 0.001), reflecting a more even distribution of genera.

**Figure 3.**
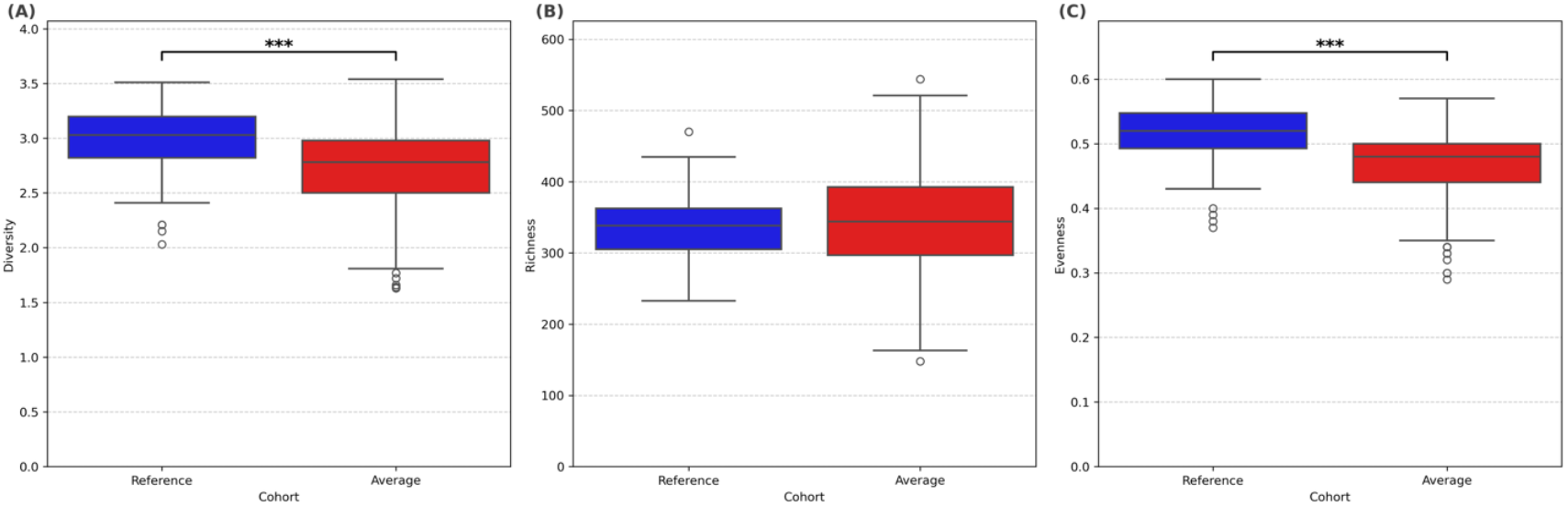
Comparison of diversity (A), richness (B), and evenness (C) between the ‘Reference’ and ‘Average’ cohorts. The box plots display the median (central line), interquartile range (box), whiskers (up to 1.5 times the interquartile range from the quartiles), and outliers (individual points). A Student’s t-test was performed to compare the means between the two cohorts. Asterisks indicate statistically significant differences (***p < 0.001) between groups for diversity (A) and evenness (C).

Together, these results suggest that the higher alpha-diversity observed in the reference cohort is driven by improved genus evenness rather than an increase in richness.

Having established differences in microbial diversity between the Reference and Average cohorts, we next investigated the functional consequences of these taxonomic shifts by analyzing the predicted metabolic capacity of the gut microbiome.

To this end, we evaluated indicators of key metabolite functions. This analysis was based on “predicted metabolism,” a metagenomic approach that infers the biochemical functions of a microbial community from its complete genetic profile. Specifically, we focused on the production capacity of short-chain fatty acids (SCFAs), such as butyrate, propionate, and acetate, which are primary end-products of non-digestible carbohydrate fermentation by the gut microbiota. We also investigated the production of vitamin B12 (cobalamin), vitamin B9 (folate) and vitamin K2. The analysis of predicted metabolic potential revealed significant differences in the capacity for producing key metabolites between the Reference and Average cohorts (Figure 4). SCFA production (Figure 4A) was significantly higher in the ‘Reference’ cohort (mean: 1450.96 ± 49 FPM) compared to the ‘Average’ cohort (mean: 1075.33 ± 15.74 FPM), with a t-test indicating a highly significant difference (***p < 0.001). Similarly, the predicted capacity for Vitamin B12 synthesis (Figure 4B) was significantly greater in the ‘Reference’ cohort (mean: 1097.68 ± 29.43 FPM) compared to the ‘Average’ cohort (mean: 848.72 ± 13.18 FPM), also showing a highly significant difference (***p < 0.001). Furthermore, Vitamin B9 (folate) (Figure 4C) production capacity followed the same trend, with the ‘Reference’ cohort exhibiting a significantly higher predicted capacity (mean: 1503.36 ± 26.51 FPM) than the ‘Average’ cohort (mean: 1389.19 ± 10.11 FPM), confirmed by a highly significant t-test result (***p < 0.001). In contrast, Vitamin K2 (Figure 4D) production capacity was significantly higher in Average cohort (mean: 910.03 ± 9.99 FPM) than in the Reference cohort (mean: 835.57 ± 24.11 FPM) (**p<0.01). These findings collectively indicate that the ‘Reference’ cohort’s microbiome possesses a superior predicted metabolic potential for producing most of these crucial host-beneficial compounds.

**Figure 4.**
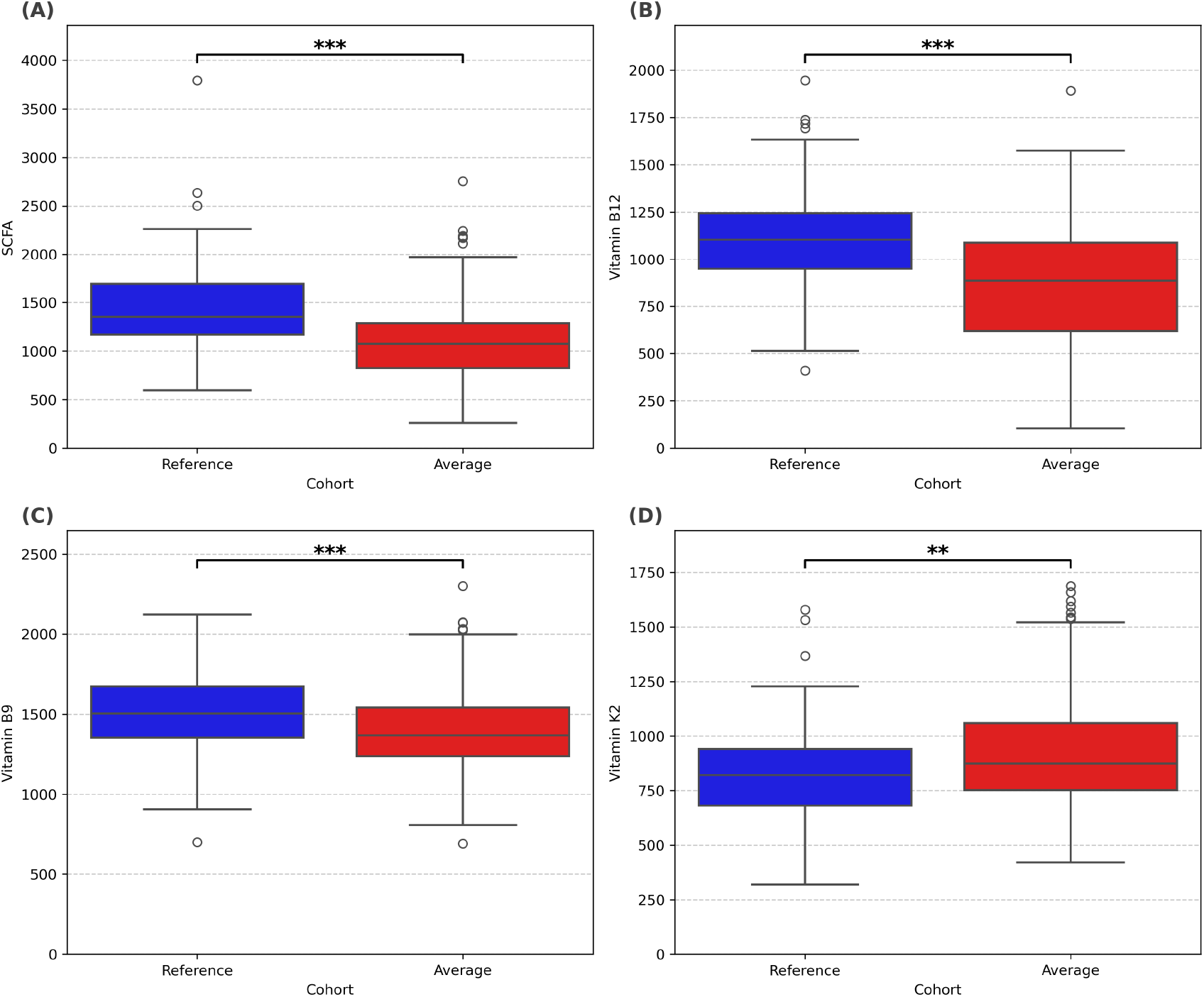
Comparison of SCFA (A), Vitamin B12 (B), Vitamin B9 (C), and Vitamin K2 (D) production capacity between the ‘Reference’ and ‘Average’ cohorts. The box plots show the fragments per million (FPM) of KEGG IDs related to the biosynthesis of each metabolite: central line indicates the median, boxes represent the interquartile range, whiskers extend up to 1.5 times the IQR from the quartiles, and individual points indicate outliers. FPM values reflect the normalized abundance of all KEGG IDs detected per sample that are involved in the metabolic pathways for each compound. A Student’s t-test was performed to compare group means. Asterisks indicate statistically significant differences (**p < 0.01; ***p < 0.001).

In addition, we evaluated specific indicators of predicted metabolic capacity related to TMA, and the synthesis of vitamins B1 (Thiamine) and B7 (Biotin), as shown in Supplementary Figure 1. The analysis revealed no significant differences in the predicted production capacities of TMA, vitamin B1, or vitamin B7 between the Reference and Average cohorts (Supplementary Figure 1A, 1B, and 1C).

### Gut microbiome variations in the average urban cohort before and after lifestyle adjustments

To investigate the impact of dietary and lifestyle changes on gut microbiome composition, we analyzed data selected from nine random participants (US01-US09) of the average urban cohort that completed two microbiome profile assessments separated by a three-months period. This approach allowed for the examination of microbiome profile shifts in response to intentional lifestyle adjustments during that period. Although these changes were not controlled or parameterized, analysis of their dietary habits revealed several consistent trends. Notably, the primary dietary shifts included a marked increase in yogurt consumption, a slight rise in the intake of vegetables and whole grains, and a significant decrease in meat consumption, while the intake of other plant-based products remained relatively stable throughout the period.

Since the genetic background of the participants could be a confounding factor, specifically polymorphisms in the *Fucosyltransferase 2* (*FUT2*) gene, we analyzed the genetic background for this biomarker in the nine participants. The results showed that only one participant was a non-secretor (US03), carrying the ‘A’ allele in a homozygous state at the SNP rs492602.

### Microbiome diversity profiles

Most individuals showed either an increase or stability in alpha diversity from T0 to T1, with no statistically significant changes (p-value > 0.05) (Figure 5A). Notably, participants US01, US04, US06 and US09 demonstrated an increase, transitioning into the reference range (indicated by the shaded green region) at T1. US08 increased alpha diversity but is still under the reference range. Conversely, US02, US03, US05 and US07 experienced a slight decrease but are still within the reference range. Overall, alpha diversity showed a general improvement, suggesting increased variety within the gut microbiome for most participants but not statistically significant.

**Figure 5.**
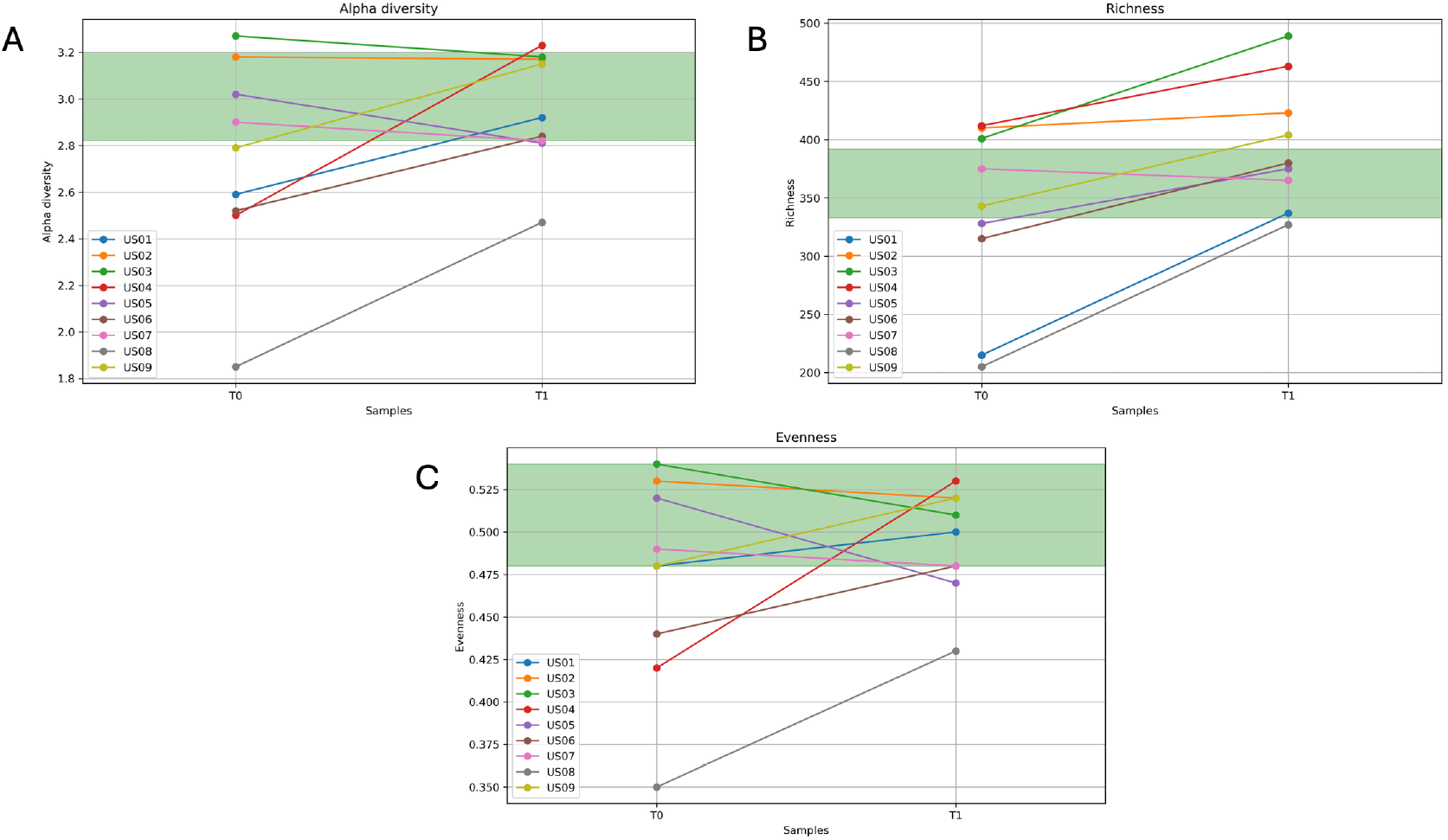
Comparison of Alpha diversity (A), genus richness (B) and genus evenness (C) of each user before (T0) and after (T1) habits improvement. Each point represents the value of each gut microbiome indicator for a user before and after habits improvements. The green zone in the graphs represents the average range observed in the reference population.

Most participants exhibited an increase in richness from T0 to T1, with some individuals (e.g., US01, US06, US08) reaching or approaching the target reference range (Figure 5B). However, US05 displayed minimal or no change, highlighting individual variability in response. The overall trend suggests a positive shift in microbial diversity, potentially influenced by the intervention applied between T0 and T1. Overall, microbial richness significantly improved between T0 and T1 (p-value: 0.0030), indicating a greater number of different microbial species post-habit changes.

Evenness demonstrated more heterogeneous responses. Several individuals (e.g., US04, US07, US09) experienced an increase in evenness, while others (e.g., US03, US05) showed a decline over the same period (Figure 5C). Many participants remained within or approached the target reference range (shaded green area in Figure 5) at T1, suggesting some convergence toward an optimal balance in the distribution of microbial taxa. However, a paired t-test comparing evenness values at T0 and T1 did not reveal statistically significant differences (p-value > 0.05). Overall, evenness showed variable changes, with no consistent trend observed across all participants.

### Metabolic capacity Profiles

SCFAs generally increased from T0 to T1 for most participants (Figure 6A). Participant US02 exhibited a particularly marked increase in SCFA production capacity, reaching the highest concentration at T1. Participants US01, US03, US04, US06, US08, and US09 also showed elevated SCFA potential production capacity, with many falling within the reference range at T1. Overall, SCFAs exhibited a significant increase in potential production capacity following the changes in habits (p-value: 0.0191).

**Figure 6.**
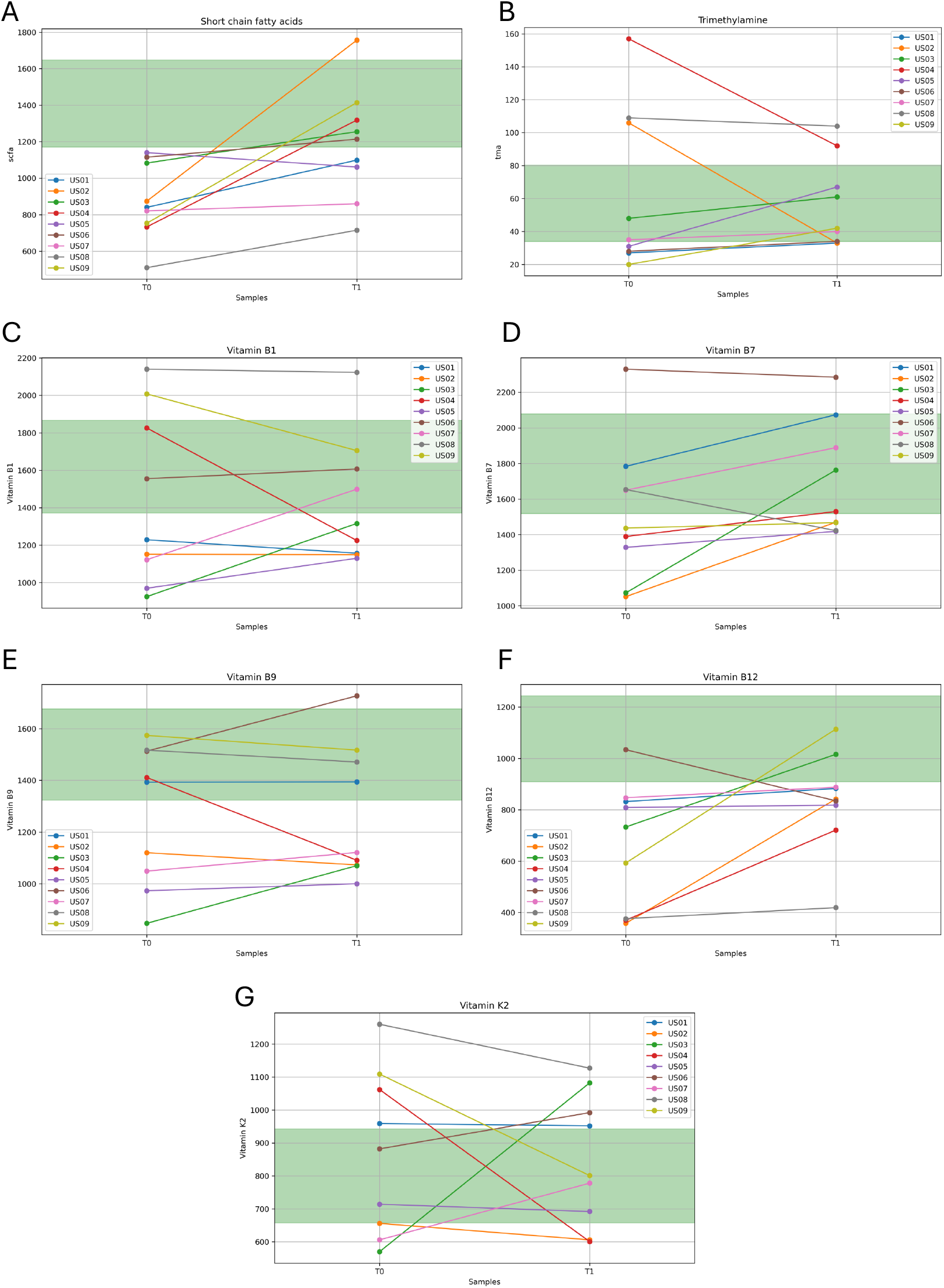
Comparison of Short chain fatty acids (A), Trimethylamine (B), Vitamin B1 (C), Vitamin B7 (D), Vitamin B9 (E), Vitamin B12 (F) and Vitamin K2 (G) production capacity of each user before (T0) and after (T1) habits improvement. Y-axis shows the fragments per million (FPM) of KEGG IDs related to the biosynthesis of each metabolite. Each point represents the value of each gut microbiome indicator for a user before and after habits improvements. The green zone in the graphs represents the average range observed in the reference population.

On the other hand, the analysis of TMA levels across two time points (T0 and T1) revealed considerable individual variability (Figure 6B). While some individuals (US02, US04, US08) showed a marked reduction in TMA levels, moving closer to or within the target reference range, others (US03, US09) experienced a slight increase. Additionally, several participants (US05, US07) exhibited relatively stable levels over time. A paired t-test comparing TMA levels at T0 and T1 yielded a p-value of 0.6386, indicating that the overall changes were not statistically significant. These results suggest that habit changes had inconsistent effects on TMA levels across the cohort, emphasizing the importance of considering individual responses when evaluating metabolic shifts.

The analysis of Vitamin B1 production capacity across two time points (T0 and T1) revealed a heterogeneous pattern among individuals (Figure 6C). Some participants (e.g., US03, US07) displayed an increase in Vitamin B1 levels, moving closer to the target reference range (shaded green area in Figure 6C), while others (e.g., US04, US09) exhibited a marked decrease. Several individuals (e.g., US01, US02, US08) maintained relatively stable levels throughout the observation period. A paired t-test comparing Vitamin B1 levels between T0 and T1 indicated no statistically significant differences in the overall cohort (p-value of 0.9868). The analysis of Vitamin B7 production capacity across T0 and T1 shows a general trend of increase for most individuals (Figure 6D). Participants such as US01, US03, and US07 exhibited significant growth, with their values moving into or closer to the target reference range (shaded green area). Conversely, some individuals, like US06, maintained stable levels, while others (e.g., US09) displayed minor increases. However, despite the changes in habits, the analysis did not reveal a statistically significant difference (p-value: 0.0803). The Vitamin B9 production capacity shows a mixed pattern (Figure 6E), with several individuals (e.g., US06, US07) demonstrating an increase in levels, while others (e.g., US04, US09) experienced a decrease. Some participants (e.g., US01, US02) remained stable throughout the study period, with their values residing within the target reference range (shaded green area). A paired t-test comparing Vitamin B9 levels at T0 and T1 resulted in a p-value of 0.8940, indicating no significant changes across the cohort. In the same way, Vitamin B12 production capacity demonstrates a noticeable increase in some individuals (Figure 6F), such as US03 and US09, with their values approaching the target reference range (shaded green area). Conversely, other participants, such as US06 and US04, exhibited stable or decreasing levels. Most individuals maintained values below the reference range despite the habit changes. A paired t-test comparing Vitamin B12 levels between T0 and T1 yielded a p-value of 0.0623. In this cohort, The Vitamin K2 production capacity shows a mixed response across participants (Figure 6G). While some individuals, such as US03 and US07, exhibited increases over time, others, including US09 and US04, showed declines. Most of the participants had values either within or close to the target reference range (shaded green area). A paired t-test indicated no statistically significant changes in the cohort (p-value: 0.8295). These findings suggest that the observed changes are highly individual and that the habits modifications did not consistently affect Vitamin B-complex and Vitamin K2 production capacity.

## DISCUSSION

In this study, we successfully reconstructed the dataset of the gut microbiome of an Argentine urban population using low-pass whole metagenome sequencing (WMS), reaffirming the method’s superiority over 16S rRNA sequencing for generating functionally relevant insights. We highlight that in a dense microbial ecosystem characterized by frequent horizontal gene transfer, understanding the collective metabolic potential is more critical than a simple taxonomic census. The capacity of low-pass WMS to move beyond taxonomy allowed us to correct for known 16S biases and, more importantly, to access the predicted functional capabilities of the microbial community.

The clustering analysis performed in the reference dataset, using the low-pass WMS technology identified three distinct cluster types dominated by the genera *Faecalibacterium, Bacteroides*, and *Prevotella*, which aligns with established ecological principles of the gut microbiome modulated by long-term dietary patterns [21].

As populations move from traditional hunter-gatherer lifestyles to industrialized urban living, there is a corresponding decline in microbial diversity and a shift in key taxa, driven primarily by a dramatic reduction in dietary fiber intake[10]. The Argentine ‘Reference’ cohort, while representing a reference urban ideal, still exhibits a microbiome profile that is substantially different from traditional populations, and the ‘Average’ cohort has moved even further along this path toward a less healthy state typical of Western societies. The concept of an “evolutionary mismatch” is a powerful explanatory framework for the observed deviation in the gut microbiome of modern western living. This concept posits that many chronic non-communicable diseases arise because the holobiont, which were shaped by thousands of years of evolution in a particular environment, are now poorly adapted to the novel environments of industrialized societies that are less than 200 years old [10], [11]. The microbiome profile of the ‘Average’ Argentine cohort is a direct biological manifestation of this profound mismatch. The modern urban diet is fundamentally deficient in the diverse array of microbiota-accessible carbohydrates (MACs), the complex plant fibers that are the preferred fuel for a robust gut ecosystem[22]. Instead, the modern diet is dominated by ultra-processed foods (UPFs), which are characterized by high loads of refined sugar, saturated and trans fats, salt, and other industrial food additives, such as emulsifiers [9].

Alpha-diversity, often quantified by indices such as the Shannon index, provides a comprehensive measure of microbial complexity within a sample, accounting for both the number of distinct taxa (richness) and the uniformity of their distribution (evenness). A higher alpha-diversity is generally associated with a more resilient and functionally robust microbiome, capable of performing a wider array of metabolic functions beneficial to the host and resisting perturbations. Genus richness refers to the total count of unique microbial genera present in a community, reflecting the sheer variety of microorganisms. While a greater number of distinct taxa can indicate a more diverse ecosystem, richness alone does not capture the relative abundances of these taxa. Finally, genus evenness quantifies how equitably the abundances of different genera are distributed within the community. High evenness suggests a balanced ecosystem where no single or few genera overwhelmingly dominate, which is often indicative of a healthy and stable microbiome. An interesting observation from the analysis of the Argentine cohorts is that the ‘Average’ urban cohort exhibits significantly lower alpha-diversity, as measured by the Shannon index, when compared to the ‘Reference’ cohort. Critically, the primary driver of this reduced diversity is not a deficit in genus richness, but a marked reduction in genus evenness. This distinction is significant because it suggests a destabilization of the community structure.

This dramatic dietary shift from our ancestors has two primary consequences for the gut microbiome. First, it effectively starves the vast communities of specialist microbes that depend on complex MACs for their survival. Without their necessary substrates, their populations decrease, leading to the loss of diversity and evenness as we observed in the Argentine ‘Average’ cohort. Second, it creates a new ecological niche that favors a different set of microbes. Organisms that are adept at thriving in a low-fiber, high-fat and high-sugar environment begin to dominate. As previously noted, the family Bacteroidaceae is a prime example of a group that is enriched in urban populations, partly due to its ability to tolerate the high concentrations of bile acids produced in response to a high-fat diet [10]. The state seen in the ‘Average’ cohort is, therefore, a predictable ecological response to a radically altered nutritional landscape.

The transition from a diverse, ancestral-type microbiome to a less diverse urban profile is reflected in the shift in the functional capacity of the entire ecosystem. The low-pass WMS approach utilized in this study allows for a crucial leap from taxonomic profiling to functional prediction. The analysis of the predicted metabolic potential of the ‘Reference’ and ‘Average’ cohorts reveals a contrast in their ability to produce key molecules that mediate host-microbe symbiosis. A significant finding from this study is the markedly lower predicted production capacity for short-chain fatty acids (SCFAs) in the ‘Average’ cohort relative to the ‘Reference’ group. This observation is the direct functional consequence of a diet deficient in the fermentable fibers (MACs) that serve as the primary substrate for SCFA-producing bacteria [23]. The predicted deficit in the ‘Average’ cohort’s ability to generate these compounds signifies a breakdown in this crucial symbiotic collaboration, with far-reaching consequences, SCFAs exert profound immunomodulatory effects by acting as ligands for a class of G-protein-coupled receptors (GPCRs), which are expressed on the surface of intestinal epithelial cells and various immune cells. The activation of these receptors by SCFAs initiates signaling cascades that are predominantly anti-inflammatory [24]. Therefore, the lower predicted SCFA production capacity in the ‘Average’ Argentine cohort could suggest a chronically weakened anti-inflammatory signaling in the gut. While the ‘Average’ cohort microbiome is characterized by a deficit in beneficial anti-inflammatory functions, the Western dietary pattern also promotes pathways that generate inflammatory environments.

TMA is generated by certain gut bacteria through the metabolism of dietary precursors such as choline, phosphatidylcholine, and L-carnitine. These precursors are particularly abundant in foods that are hallmarks of the Western diet, including red meat, processed meats, eggs, and high-fat dairy products. Once produced in the gut, TMA is absorbed into the bloodstream, transported to the liver, and rapidly oxidized by the enzyme flavin-containing monooxygenase 3 (FMO3) into trimethylamine N-oxide (TMAO) [25]. A large and growing body of evidence has established TMAO as a contributor to the pathogenesis of cardiovascular disease (CVD). Elevated plasma levels of TMAO have been shown in numerous clinical studies to be a strong, independent predictor of major adverse cardiovascular events, including heart attack, stroke, and all-cause mortality, even after adjusting for traditional risk factors [26].

The gut microbiome plays a pivotal role in human health, extending its influence on the production of various B-complex vitamins, such as B1 (thiamine), B7 (biotin), B9 (folate), and B12 (cobalamin), as well as vitamin K2 (menaquinone) [27]. These vitamins are indispensable cofactors for a wide array of host biological processes, including energy metabolism, DNA synthesis and repair, nervous system function, blood coagulation, and bone health [28]. Given the limited or absent *de novo* synthesis capacity of these vitamins in human cells, their microbial production, primarily in the distal gut from unabsorbed dietary components like fermentable fibers, is crucial for maintaining adequate vitamin status and overall physiological homeostasis [28]. Our analysis showed that the ‘Reference’ cohort exhibited a significantly higher predicted capacity for the synthesis of Vitamin B9, B12 and K2 compared to the ‘Average’ urban population cohort, while no statistically significant differences were observed for Vitamin B1 or B7 between the cohorts. These observed deficits in the predicted vitamin synthesis capacity in the ‘Average’ urban cohort align with the broader understanding of how modern, industrialized diets, often low in fermentable fibers and high in ultra-processed foods, negatively impact the gut microbiome [10]. This functional compromise highlights the mentioned “evolutionary mismatch” between ancestral human biology and contemporary dietary environments. The reduced availability of microbiota-accessible carbohydrates (MACs) in modern diets effectively starves beneficial, specialized microbes, leading to a reduction in functional diversity and compromised metabolic potential. While our study effectively assessed the predicted capacity for vitamin synthesis, it is crucial to acknowledge that this does not always equate to the actual bioavailability or *contribution* to the host’s vitamin status. Future research should delve into the specific forms of vitamins produced and their precise bioavailability to the host to fully elucidate their physiological impact.

The longitudinal analysis of participants who implemented lifestyle changes provides a crucial proof-of-concept for the plasticity of the gut microbiome. We observed statistically significant improvements in microbial richness and, most importantly, in the predicted capacity for SCFA production, demonstrating that positive modulation is achievable. However, the results also highlighted a remarkable degree of inter-individual variability. Responses in TMA levels and the synthesis of several B vitamins and vitamin K2 were highly personalized, with no consistent trend across the cohort. This “responder” versus “non-responder” phenomenon is a well-documented issue in the field of clinical nutrition and microbiome research [29]. This finding challenges the efficacy of a universal, one-size-fits-all approach to dietary and lifestyle recommendations, suggesting that such strategies are destined to be suboptimal for a significant fraction of any given population. The ‘average’ gut microbiome is a collection of highly diverse, individualized ecosystems, each possessing a different capacity to adapt and respond to change.

Arguably, one of the most important determinants of how an individual gut ecosystem responds to a dietary change is the composition and functional capacity of their microbiome at baseline. The gut microbiome can be viewed as a vast enzymatic factory. The ability to break down a specific type of dietary fiber and convert it into beneficial metabolites like butyrate is entirely contingent on possessing the requisite microbial species that harbor the necessary carbohydrate-active enzymes (CAZymes) and other enzymes [30]. If an individual’s baseline microbiome, perhaps because of long-term dietary patterns or antibiotic use, has lost the key bacterial species specialized in fermenting a particular fiber (e.g., inulin, pectin, or resistant starch), then supplementing with that fiber will be ineffective [31]. The varied responses in the Argentine urban-uncontrolled cohort are likely a direct reflection of the differences in their starting microbial community structures. Some individuals likely retained a sufficient diversity of fiber-degraders to be “responders,” while others may have crossed a threshold of microbial depletion, rendering them “non-responders” to the specific dietary changes.

While diet and other environmental factors are the dominant forces shaping the gut microbiome, host genetics also exert a significant—albeit more subtle—influence. Genetic variations in the host can affect numerous aspects of the gut environment, thereby shaping the microbial communities that can thrive there. These include genes that control the immune system’s interaction with microbes (e.g., pattern recognition and inflammatory response genes), genes involved in the synthesis and composition of the gut’s protective mucus layer (which serves both as a barrier and a food source for certain microbes), and genes that regulate host metabolism [32]. This genetic background creates a unique “host filter” that contributes to the high degree of inter-individual variability in baseline microbiome composition and, by extension, in responses to dietary interventions. In our study, we analyzed the genetic background of the nine participants, focusing on the *FUT2* gene. A non-functional *FUT2* gene results in the absence of fucosylated glycans (HBGAs) on mucosal surfaces and in secretions. This has important downstream consequences for non-secretors, particularly in terms of host–microbe interactions and nutrient processing. Without fucosylated glycans, non-secretors lack a critical nutrient source and adhesion site for beneficial bacteria, which can lead to reduced microbial diversity and metabolic capacity [33]. In our cohort, only one out of nine participants was a non-secretor of HBGAs. This finding suggests that while most participants possess normal fucosylated glycan secretion, their individual responses to dietary interventions can still vary widely, likely influenced by other host and microbial factors.

In conclusion, our study validated the use of low-pass WMS to uncover the potential metabolic capacity and functional deficits in the gut microbiome of an urban population. The significant heterogeneity in response to lifestyle intervention strongly suggests that universal dietary recommendations are insufficient. This variability is likely governed by a combination of factors, including an individual baseline microbiome composition, host genetics, and the specific nature of the dietary changes implemented. Therefore, our findings support a paradigm shift towards personalized nutritional strategies. By leveraging functional microbiome profiling to understand individual metabolic potential and deficiencies, it becomes possible to design targeted interventions aimed at restoring a more balanced and functionally robust gut ecosystem, thereby mitigating the long-term health risks associated with the evolutionary mismatch of the modern diet.

Finally, we acknowledge some limitations. Although low-pass WMS provides superior resolution compared to 16S rRNA sequencing, the predicted metabolic capacities do not necessarily reflect the actual bioavailability of metabolites. Future studies should incorporate metabolomics to validate these findings. Additionally, the small sample size of the longitudinal analysis (n=9) limits generalizability, and larger studies are needed to confirm the plasticity of the gut microbiome in these urban populations. Additionally, exploring factors such as host genetics in depth and the interaction with the gut microbiome could shed light on the observed interindividual variability.

## Acknowledgements

The authors want to thank all members of the lab and the Heritas team, specifically Sofia Sidlik, Sofia Collazo, Florencia Riccomi, Bianca Brun for helpful discussions during the preparation of this manuscript, to all Rewell participants for contributing their data anonymously, to the nutrition health coaches Carla Marcuzzi, Surya Perez for their knowledge, patience and contributions.

## Data and Code availability

<u>militrot/reference_database</u>. Dataset availability after final publication.

## Supplemental Material

**Supplementary Figure 1.**
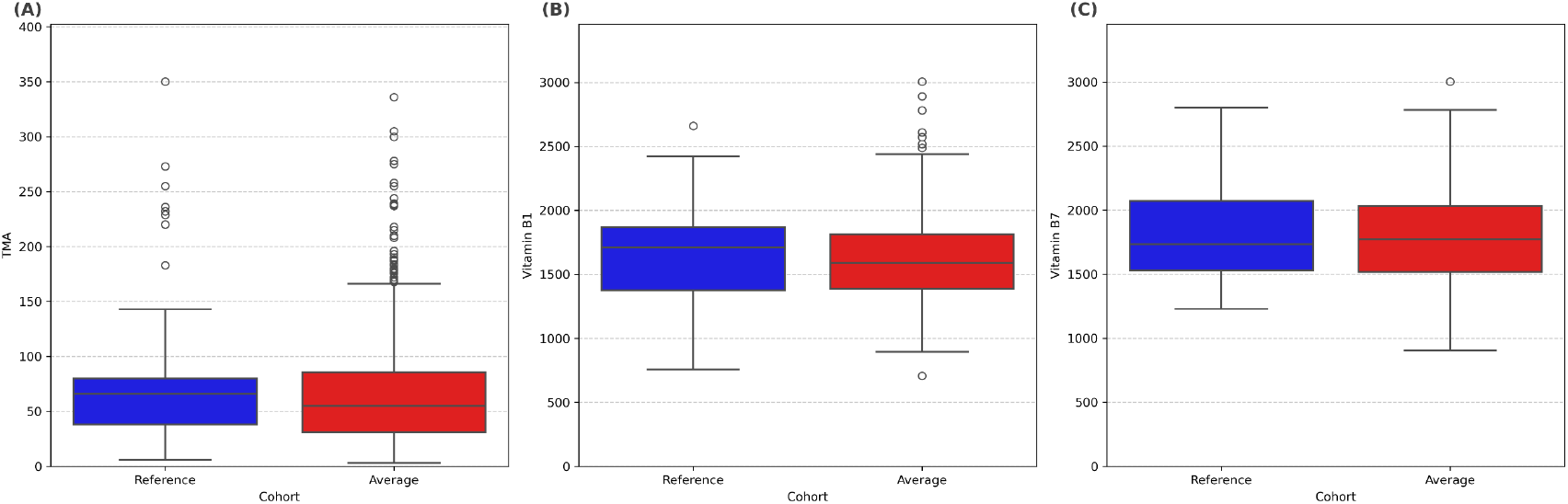
Comparison of TMA (A), Vitamin B1 (B) and Vitamin B7 (C) production capacity between the ‘Reference’ and ‘Average’ cohorts. The box plots show the fragments per million (FPM) of KEGG IDs related to the biosynthesis of each metabolite. Each box plots display the median (central line), interquartile range (box), whiskers (up to 1.5 times the interquartile range from the quartiles), and outliers (individual points). A Student’s t-test was performed to compare the means between the two cohorts. Asterisks indicate statistically significant differences (*p < 0.05) between groups.

***Supplementary file 1***. *See separate pdf file*

***Supplementary table 1***. See separate excel file

## REFERENCES

[1] A. B. Shreiner, J. Y. Kao, and V. B. Young, ‘The gut microbiome in health and in disease’, Curr Opin Gastroenterol, vol. 31, no. 1, pp. 69–75, Jan. 2015, doi: 10.1097/MOG.0000000000000139,.

[2] C. Foppa, T. Rizkala, A. Repici, C. Hassan, and A. Spinelli, ‘Microbiota and IBD: Current knowledge and future perspectives’, Digestive and Liver Disease, vol. 56, no. 6, pp. 911–922, Jun. 2024, doi: 10.1016/j.dld.2023.11.015.

[3] K.-R. Baek, S. Singh, H.-S. Hwang, and S.-O. Seo, ‘Using Gut Microbiota Modulation as a Precision Strategy Against Obesity’, Int J Mol Sci, vol. 26, no. 13, p. 6282, Jun. 2025, doi: 10.3390/IJMS26136282.

[4] J. Wang et al., ‘A metagenome-wide association study of gut microbiota in type 2 diabetes’, Nature, vol. 490, no. 7418, pp. 55–60, Oct. 2012, doi: 10.1038/NATURE11450,.

[5] T. Katsimichas, P. Theofilis, K. Tsioufis, and D. Tousoulis, ‘Gut Microbiota and Coronary Artery Disease: Current Therapeutic Perspectives’, Metabolites, vol. 13, no. 2, p. 256, Feb. 2023, doi: 10.3390/METABO13020256.

[6] M. M. Nakhal et al., ‘The Microbiota–Gut–Brain Axis and Neurological Disorders: A Comprehensive Review’, Life, vol. 14, no. 10, p. 1234, Oct. 2024, doi: 10.3390/LIFE14101234.

[7] E. J. Song and J. H. Shin, ‘Personalized Diets based on the Gut Microbiome as a Target for Health Maintenance: from Current Evidence to Future Possibilities’, J Microbiol Biotechnol, vol. 32, no. 12, p. 1497, Dec. 2022, doi: 10.4014/JMB.2209.09050.

[8] S. Gurunathan, P. Thangaraj, and J. H. Kim, ‘Postbiotics: Functional Food Materials and Therapeutic Agents for Cancer, Diabetes, and Inflammatory Diseases’, Foods, vol. 13, no. 1, Jan. 2024, doi: 10.3390/FOODS13010089,.

[9] C. A. Monteiro et al., ‘Ultra-processed foods: What they are and how to identify them’, Public Health Nutr, vol. 22, no. 5, pp. 936–941, Apr. 2019, doi: 10.1017/S1368980018003762,.

[10] S. Rosas-Plaza, A. Hernández-Terán, M. Navarro-Díaz, A. E. Escalante, R. Morales-Espinosa, and R. Cerritos, ‘Human Gut Microbiome Across Different Lifestyles: From Hunter-Gatherers to Urban Populations’, Front Microbiol, vol. 13, Apr. 2022, doi: 10.3389/FMICB.2022.843170/PDF.

[11] M. K. Zinöcker and I. A. Lindseth, ‘The western diet–microbiome-host interaction and its role in metabolic disease’, Nutrients, vol. 10, no. 3, Mar. 2018, doi: 10.3390/NU10030365,.

[12] H. Tilg and A. R. Moschen, ‘Food, immunity, and the microbiome’, Gastroenterology, vol. 148, no. 6, pp. 1107–1119, May 2015, doi: 10.1053/j.gastro.2014.12.036.

[13] D. Rondinella et al., ‘The Detrimental Impact of Ultra-Processed Foods on the Human Gut Microbiome and Gut Barrier’, Nutrients, vol. 17, no. 5, p. 859, Mar. 2025, doi: 10.3390/NU17050859.

[14] J. Jovel et al., ‘Characterization of the Gut Microbiome Using 16S or Shotgun Metagenomics’, Front Microbiol, vol. 7, no. APR, p. 459, Apr. 2016, doi: 10.3389/FMICB.2016.00459.

[15] I. Laudadio, V. Fulci, F. Palone, L. Stronati, S. Cucchiara, and C. Carissimi, ‘Quantitative Assessment of Shotgun Metagenomics and 16S rDNA Amplicon Sequencing in the Study of Human Gut Microbiome’, OMICS, vol. 22, no. 4, pp. 248–254, Apr. 2018, doi: 10.1089/OMI.2018.0013,.

[16] S. Rezasoltani, D. A. Bashirzadeh, E. N. Mojarad, H. A. Aghdaei, M. Norouzinia, and S. Shahrokh, ‘Signature of gut microbiome by conventional and advanced analysis techniques: Advantages and disadvantages’, Middle East J Dig Dis, vol. 12, no. 1, pp. 249–255, Jan. 2020, doi: 10.15171/MEJDD.2020.157,.

[17] R. Ranjan, A. Rani, A. Metwally, H. S. McGee, and D. L. Perkins, ‘Analysis of the microbiome: Advantages of whole genome shotgun versus 16S amplicon sequencing’, Biochem Biophys Res Commun, vol. 469, no. 4, pp. 967–977, Jan. 2016, doi: 10.1016/j.bbrc.2015.12.083.

[18] R. Hajjo, D. A. Sabbah, and A. Q. Al Bawab, ‘Unlocking the Potential of the Human Microbiome for Identifying Disease Diagnostic Biomarkers’, Diagnostics, vol. 12, no. 7, p. 1742, Jul. 2022, doi: 10.3390/DIAGNOSTICS12071742.

[19] C. Rohr, M. Sciara, B. Brun, F. Fay, and M. P. Vazquez, ‘Generation of a robust reference gut microbiome dataset for an urban population in Argentina optimized by a machine learning approach’, bioRxiv, p. 2023.06.24.546376, Jun. 2023, doi: 10.1101/2023.06.24.546376.

[20] J. Tamames and F. Puente-Sánchez, ‘SqueezeMeta, a highly portable, fully automatic metagenomic analysis pipeline’, Front Microbiol, vol. 10, no. JAN, 2019, doi: 10.3389/FMICB.2018.03349,.

[21] P. I. Costea et al., ‘Enterotypes in the landscape of gut microbial community composition’, Nat Microbiol, vol. 3, no. 1, p. 8, 2017, doi: 10.1038/S41564-017-0072-8.

[22] E. D. Sonnenburg, S. A. Smits, M. Tikhonov, S. K. Higginbottom, N. S. Wingreen, and J. L. Sonnenburg, ‘Diet-induced extinction in the gut microbiota compounds over generations’, Nature, vol. 529, no. 7585, p. 212, Jan. 2016, doi: 10.1038/NATURE16504.

[23] G. Ayakdaş and D. Ağagündüz, ‘Microbiota-accessible carbohydrates (MACs) as novel gut microbiome modulators in noncommunicable diseases’, Heliyon, vol. 9, no. 9, p. e19888, Sep. 2023, doi: 10.1016/J.HELIYON.2023.E19888.

[24] S. Facchin et al., ‘Short-Chain Fatty Acids and Human Health: From Metabolic Pathways to Current Therapeutic Implications’, Life 2024, Vol. 14, Page 559, vol. 14, no. 5, p. 559, Apr. 2024, doi: 10.3390/LIFE14050559.

[25] Y. Zhang, Y. Wang, B. Ke, and J. Du, ‘TMAO: how gut microbiota contributes to heart failure’, Translational Research, vol. 228, pp. 109–125, Feb. 2021, doi: 10.1016/J.TRSL.2020.08.007.

[26] J. Zhen et al., ‘The gut microbial metabolite trimethylamine N-oxide and cardiovascular diseases’, Front Endocrinol (Lausanne), vol. 14, p. 1085041, Feb. 2023, doi: 10.3389/FENDO.2023.1085041/FULL.

[27] C. Tarracchini et al., ‘Exploring the vitamin biosynthesis landscape of the human gut microbiota’, mSystems, vol. 9, no. 10, Oct. 2024, doi: 10.1128/MSYSTEMS.00929-24/SUPPL_FILE/MSYSTEMS.00929-24-S0003.XLSX.

[28] S. Magnúsdóttir, D. Ravcheev, V. De Crécy-Lagard, and I. Thiele, ‘Systematic genome assessment of B-vitamin biosynthesis suggests cooperation among gut microbes’, Front Genet, vol. 6, no. MAR, p. 129714, Apr. 2015, doi: 10.3389/FGENE.2015.00148/BIBTEX.

[29] C. R. Kok, D. Rose, and R. Hutkins, ‘Predicting Personalized Responses to Dietary Fiber Interventions: Opportunities for Modulation of the Gut Microbiome to Improve Health’, Annu Rev Food Sci Technol, vol. 14, pp. 157–182, Mar. 2023, doi: 10.1146/ANNUREV-FOOD-060721-015516,.

[30] J. F. Wardman, R. K. Bains, P. Rahfeld, and S. G. Withers, ‘Carbohydrate-active enzymes (CAZymes) in the gut microbiome’, Nat Rev Microbiol, vol. 20, no. 9, pp. 542–556, Sep. 2022, doi: 10.1038/S41579-022-00712-1,.

[31] S. O. Onyango, J. Juma, K. De Paepe, and T. Van de Wiele, ‘Oral and Gut Microbial Carbohydrate-Active Enzymes Landscape in Health and Disease’, Front Microbiol, vol. 12, Dec. 2021, doi: 10.3389/FMICB.2021.653448,.

[32] A. Horowitz, S. D. Chanez-Paredes, X. Haest, and J. R. Turner, ‘Paracellular permeability and tight junction regulation in gut health and disease’, Nature Reviews Gastroenterology & Hepatology 2023 20:7, vol. 20, no. 7, pp. 417–432, Apr. 2023, doi: 10.1038/s41575-023-00766-3.

[33] O. Giampaoli, G. Conta, R. Calvani, and A. Miccheli, ‘Can the FUT2 Non-secretor Phenotype Associated With Gut Microbiota Increase the Children Susceptibility for Type 1 Diabetes? A Mini Review’, Front Nutr, vol. 7, p. 606171, Dec. 2020, doi: 10.3389/FNUT.2020.606171/FULL.

