## Supplementary File 1 for "Using low pass whole metagenome sequencing for gut microbiome profiling in an Argentine urban population"

**SECTION 1: PERSONAL DATA**

1. Indicate your current weight in kilograms (kg):
2. Indicate the contour or circumference of your waist at navel height in centimeters (cm):

**Do you currently smoke any tobacco products or derivatives such as cigarettes, cigars, or a pipe?**

- o YES
- o NO

3. On average, how many cigarettes do you smoke per day?
4. **Are you currently pregnant?**
   - o YES
   - o NO
5. **Do you take or have you taken any type of medication in the last month? If so, please select the option(s) as appropriate:**
   - o I have not taken any medication in the last month, nor do I take chronic medications.
   - o Antiallergics (such as Loratadine, among others)
   - o Antibiotics (of any type)
   - o Contraceptives
   - o Antivirals (such as acyclovir, among others)
   - o Aspirin
   - o Corticosteroids
   - o Analgesics (paracetamol)
   - o Nonsteroidal anti-inflammatory drugs / NSAIDs (Ibuprofen, diclofenac)
   - o Levothyroxine
   - o Chemotherapy medication
   - o Medications for stomach reflux (such as Omeprazole and others)
   - o Medications for diabetes or blood sugar such as metformin
   - o Medications for heart health such as: Enalapril, Rosuvastatin, Statins
   - o Medications to help sleep
   - o Medications for mental health
   - o Antacids

- Other/s

6. **In a typical week, which of these vitamins and/or supplements do you take most days? Please select all that apply:**
    - Multivitamins
    - Melatonin
    - Omega 3
    - Vitamin A
    - Vitamin B1-B6
    - Biotin
    - Folic acid
    - Vitamin C
    - Vitamin D
    - Vitamin E, H, or K
    - Calcium, magnesium, iron, zinc, selenium, or potassium
    - Probiotics
    - Prebiotics
    - Collagen
    - Other, specify which:
7. **What is your bowel movement frequency?**
    - One or more times a day
    - More than 3 times a week
    - Less than 3 times a week
8. **The Bristol stool scale is used in medicine to classify the shape of stools into 7 groups. Often, this gives us an indication of your intestinal health and transit time in your intestine. We are interested in knowing, in your case, what the shape of your stools is normally. Please select an option below:**
    - Type 1: Separate hard lumps, like nuts
    - Type 2: Sausage-shaped but lumpy
    - Type 3: Like a sausage but with cracks on its surface
    - Type 4: Like a sausage or snake, smooth and soft
    - Type 5: Soft blobs with clear-cut edges
    - Type 6: Fluffy pieces with ragged edges, a mushy stool

- Type 7: Watery, no solid pieces, entirely liquid

**SECTION 3: PHYSICAL ACTIVITY**

9. **How many hours a day do you spend sitting or lying down? (On average, excluding sleeping hours).**
   - 4 hours or less
   - Between 4 and 8 hours
   - More than 8 hours
10. **Do you do aerobic physical exercise? By aerobic, we specifically mean that which makes you sweat or increases your breathing rate, such as brisk walking, cycling, running, etc.**
    - YES
    - NO
11. **How much time per week do you dedicate to aerobic physical exercise?**
    - 30 minutes or less per week
    - 60 minutes a week
    - 90 minutes a week
    - 120 minutes a week
    - 150 minutes or more a week
12. **Do you perform muscle-strengthening exercises such as squats, crunches, weights, yoga, pilates, etc.?**
    - YES
    - NO
13. **How often do you perform muscle-strengthening exercises such as squats, crunches, weights, yoga, pilates, etc.?**
    - 30 minutes or less per week
    - 60 minutes a week
    - 90 minutes a week
    - 120 minutes a week
    - 150 minutes or more a week

**SECTION 4: DIET**

14. **How would you define your usual type of diet?**
    - Vegan
    - Vegetarian

- Gluten-free
- Mediterranean type (Base of plant-based foods, moderate consumption of fish, dairy, and poultry. Occasional consumption of meats and sweet foods)
- Low-carb (Paleo or Ketogenic)
- Varied with foods of animal and plant origin
- Low in FODMAPS (Excludes certain dietary components such as fructans, galactans, fructose, lactose, polyols)
- I don't know, I don't follow any specific diet
- Other

15. **What meals do you usually have in a day? Mark all that apply.**
    - Breakfast
    - Lunch
    - Afternoon snack
    - Dinner
    - Snacks between meals
16. **Approximately, how many hours pass from the last intake of the day (e.g., dinner) to the first intake of the next day (e.g., breakfast)?**
    - Less than 8 hours
    - From 8 to 10 hours
    - From 10 to 12 hours
    - From 12 to 16 hours
    - 16 hours or more
17. **In what way and under what circumstances do you usually have your meals? Please select all the options you consider correct:**
    - I eat quickly because I have little time.
    - I take at least 20 minutes to eat.
    - I don't pay attention to chewing.
    - I pay attention to chewing.
    - I have access to electronic devices (cell phones, television, iPad, computer, etc.).
    - I do not have access to electronic devices (cell phones, television, computer, etc.).
    - I usually sit in a comfortable place.

- I do it standing due to lack of space or time.

18. **How many servings of vegetables do you consume on average per day? Consider all types of vegetables, both raw and cooked, consumed in salads, stews, soups, and hot side dishes (vegetables do not include potatoes, sweet potatoes, or corn). 1 serving of vegetables equals 1 cup or 1 small starter plate - 2 servings of vegetables equals 2 cups or 1 deep plate.**
    - None or less than 1 serving per day
    - 1 serving per day
    - 2 servings per day
    - 3 or more servings per day
19. **How many servings of fruit do you consume on average per day? Consider all types of fruits, both raw and cooked, fresh or dehydrated (figs, raisins, etc.). 1 serving of fruit equals 1 large unit (apple, pear, peach, orange, banana, etc.), 2 small units (kiwi, tangerines), or 1 cup of cut fruit.**
    - None or less than 1 serving per day
    - 1 serving per day
    - 2 servings per day
    - 3 or more servings per day
20. **How many servings of legumes do you consume per week? Consider legumes as chickpeas, lentils, beans, fava beans, peas. 1 serving of legumes equals 1 cup or a small plate.**
    - None or less than 1 serving a week
    - 1 serving a week
    - 2 servings a week
    - More than 2 servings a week
21. **How many servings of nuts do you consume per week? Consider nuts as peanuts, almonds, walnuts, pistachios, cashews, hazelnuts, etc. 1 serving equals a handful of nuts.**
    - None or less than 1 serving a week
    - 1 serving a week
    - 2 servings a week
    - More than 2 servings a week
22. **How many servings of whole grains do you consume per day? Whole grains are considered to be brown rice or whole wheat pasta, whole wheat bread, quinoa, oats, whole grain breakfast cereals, whole grain doughs, or dishes made with whole grains. 1 serving equals 1 cup of ready-to-serve brown rice or**

**whole wheat pasta, 1 cup of whole grain breakfast cereals, or 2 slices of whole wheat bread, 1 serving of whole grain dough.**

- None or less than 1 serving per day
- 1 serving per day
- 2 servings per day
- More than 2 servings per day
- Some servings a week

23. **In a week, how many meals with meat do you consume? Consider meats as chicken, turkey, beef, veal, pork, and their derivatives such as hamburgers, sausages, offal, and pre-made foods, among others.**
    - I don't consume
    - From 1 to 4 times a week
    - From 5 to 8 times a week
    - From 9 to 14 times a week
24. **How many times a week do you consume fish? Consider all types of fish, fresh or frozen, and canned fish.**
    - I don't consume
    - 1 time a week
    - 2 times a week
    - More than 2 times a week
25. **How many times a week do you consume yogurt? Consider yogurt made with both animal milk and vegetable drink.**
    - I don't consume
    - 1 time a week
    - 2 to 3 times a week
    - 5 or 6 times a week
26. **How often do you consume fermented foods (other than yogurt)? Example of fermented foods: kefir, kombucha, sauerkraut, among others.**
    - I don't consume
    - Sometime during the month
    - Every week
27. **How often do you consume cow's milk?**
    - I don't consume

- 1 time a week
- 2 to 3 times a week
- 5 or 6 times a week

28. **How often do you consume eggs?**
    - I don't consume
    - Occasionally, I consume some during the week
    - 1 to 2 units per day
    - More than 2 units per day
29. **How often do you consume cheese (derived from animal milk)?**
    - I don't consume
    - 1 time a day
    - More than 1 time a day
    - Occasionally, only a few times a week.
30. **What type of oil or fat do you use most often for cooking?**
    - Corn or sunflower oil
    - Olive oil
    - Canola or high-oleic sunflower oil
    - Butter or margarine
    - Coconut oil
    - Other
31. **How often do you consume seeds like chia, flax, sunflower seeds, pumpkin seeds?**
    - I don't consume
    - Occasionally
    - 2 to 3 times per week
    - 4 or more times per week
32. **How many different plant-based foods do you consume in an average week? For this question, it is important that you take a few minutes to think about the variety of plant-based foods you consume on average per week. We suggest you take notes to calculate the average for the last week. You can list the different plant-based foods you consumed at each meal last week. Include cereals (rice, oats, corn, wheat, quinoa, etc.), legumes (lentils, peas, chickpeas, fava beans, beans), vegetables, fruits, nuts (almonds, walnuts, peanuts, hazelnuts, etc.), seeds (chia, flax, sunflower, etc.).**

- Less than 10 varieties a week
- From 10 to 19 varieties a week
- From 20 to 29 varieties a week
- 30 or more varieties

33. **On average, how many times per week do you consume food that was not prepared at home?**
    - Less than 2 times a week
    - 2 to 3 times a week
    - 4 or more times a week
34. **Could you indicate your average daily consumption of water in glasses? For reference, measure in 250 ml glasses.**
35. **Could you indicate your approximate daily consumption of coffee in cups? For reference, a standard cup measure of approximately 230 ml can be used.**
36. **Could you indicate your average weekly consumption of alcoholic beverages in ml? Indicate the total for the week (in ml). For reference: 1 large can of beer is equivalent to 500 ml, 1 medium glass of wine to 175 ml, 1 large glass of wine to 250 ml, 1 bottle of wine or other beverages to 750 ml, 1 measure of spirits to 30 ml.**
37. **How often do you consume artificial drinks? Artificial drinks are considered powdered and/or concentrated juices, sodas, industrial flavored waters, etc.**
    - I don't consume
    - 1 time a week
    - 2 to 3 times per week
    - 4 to 7 times per week
38. **How often do you consume pastry products, croissants, sweet cookies, sugary cereals, etc.?**
    - I don't consume
    - 1 time a week
    - 2 to 3 times per week
    - 4 to 7 times per week
39. **What do you use to sweeten drinks and food?**
    - I don't add sweetener
    - Common, muscovado, brown, or whole sugar
    - 100% stevia sweetener

- o Honey
- o Other type of sweetener or sweetening agent

**SECTION 5 - REST AND STRESS**

40. **Regarding your current daily stress level, please indicate what best approximates it on a scale of 1 to 10, where 1 is the lowest stress for most of the day, and 10 is the maximum stress throughout the day:**
41. **How many hours of sleep on average do you get per day?**
42. **From 1 to 10, where 1 is very bad and 10 is very good, how would you rate your average sleep?**
43. **How do you feel when you wake up in the morning after sleeping?**
    - o I almost always wake up tired.
    - o Sometimes I wake up tired.
    - o I almost always wake up rested.
44. **Do you wake up during the night?**
    - o Almost every night
    - o Some nights
    - o Some nights, but due to external factors (a pet, a child, external noises)
    - o Almost never
45. **How long does it take you to fall asleep?**
    - o A few minutes, it's very fast.
    - o Between 10 to 20 minutes
    - o Between 20 to 30 minutes
    - o At least 30 minutes
    - o 1 hour or more
46. **In the 2 hours before you go to sleep, for how long do you use electronic devices? For example: reading on your cell phone, using a computer, or watching television.**
    - o I don't use them.
    - o Less than 10 minutes
    - o Between 10 to 20 minutes
    - o Between 20 to 30 minutes
    - o At least 30 minutes
    - o 1 hour or more

47. **How often do you have contact with nature? Consider activities such as walking in parks, walking barefoot on the grass, gardening, etc.**
    - At least once a week
    - At least once every 15 days
    - At least once a month
    - At least once every 6 months
    - I don't remember the last time I did this
